# Evolution of *Rhodopseudomonas palustris* to degrade halogenated aromatic compounds involves changes in pathway regulation and enzyme specificity

**DOI:** 10.1101/2023.10.10.561740

**Authors:** Irshad Ul Haq, Annika Christensen, Kathryn R. Fixen

## Abstract

Halogenated aromatic compounds are used in a variety of industrial applications but can be harmful to humans and animals when released into the environment. Microorganisms that degrade halogenated aromatic compounds anaerobically have been isolated but the evolutionary path that they may have taken to acquire this ability is not well understood. A strain of the purple nonsulfur bacterium, *Rhodopseudomonas palustris*, RCB100, can use 3-chlorobenzoate (3-CBA) as a carbon source whereas a closely related strain, CGA009, cannot. To reconstruct the evolutionary events that enabled RCB100 to degrade 3-CBA, we selected for and isolated a CGA009 strain capable of growing on 3-CBA, although not as well as RCB100. Comparative whole-genome sequencing of the evolved strain and RCB100 revealed large deletions encompassing *badM*, a transcriptional repressor of genes for anaerobic benzoate degradation. It was previously shown that in strain RCB100, a single nucleotide change in an alicyclic acid coenzyme A ligase gene, named *aliA*, gives rise to a variant AliA enzyme that has high activity with 3-CBA. When we introduced the RCB100 *aliA* allele and a *badM* deletion into *R. palustris* CGA009, it grew on 3-CBA at a similar rate as RCB100. This work provides an example of pathway evolution that includes a variant of a promiscuous enzyme with enhanced substrate specificity and a regulatory mutation that leads to constitutive expression of a pathway that does not regulate the promiscuous enzyme.

**Importance:** Biodegradation of man-made compounds often involves the activity of promiscuous enzymes whose native substrate is structurally similar to the man-made compound. Based on the enzymes involved, it is possible to predict what microorganisms are likely involved in biodegradation of anthropogenic compounds. However, there are examples of organisms that contain the required enzyme(s) and yet cannot metabolize these compounds. We found that even when the purple nonsulfur bacterium, *Rhodopseudomonas palustris*, encodes all the enzymes required for degradation of a halogenated aromatic compound, it is unable to metabolize that compound. Using adaptive evolution, we found a regulatory mutation and a variant of promiscuous enzyme with increased substrate specificity were required, but the ability to metabolize a halogenated aromatic compound also resulted in reduced fitness on another aromatic compound. This work provides insight into how an environmental isolated evolved to use halogenated aromatic compounds and the potential ecological trade-offs associated with this adaptation.

## Introduction

Halogenated aromatic compounds are widely used in a variety of applications but also have harmful effects on human and animal health (1, 2). Their widespread use has led to their accumulation in the environment, and much research has been done to understand the role microorganisms play in degrading these compounds (2). One such example is the purple nonsulfur bacterium, *Rhodopseudomonas palustris* RCB100. *R. palustris* RCB100 has been shown to metabolize 3-chlorobenzoate (3-CBA) during photoheterotrophic growth using the benzoyl-CoA degradation pathway (3). In this pathway, aromatic compounds are first activated by a coenzyme A (CoA) ligase to benzoyl-CoA and then reduced by the enzyme benzoyl-CoA reductase (Fig. 1) (4). In *R. palustris* RCB100, the CoA ligase responsible for activating 3-CBA shows a higher specificity for chlorobenzoates, and this specificity is due to a single nucleotide change in *aliA*, which encodes an alicyclic acid CoA ligase (5). This change results in a variant of AliA with a serine instead of a threonine found in other *R. palustris* strains at amino acid 208 (T208S). AliA^T208S^ has 10-fold more activity with 3-CBA than its native substrate (5). After activation by a CoA ligase, 3-chlorobenzoyl-CoA is reductively dehalogenated by benzoyl-CoA reductase, which can reductively dehalogenate a number of different halobenzoyl-CoA substrates (Fig. 1) (6–8).

**Figure 1.**
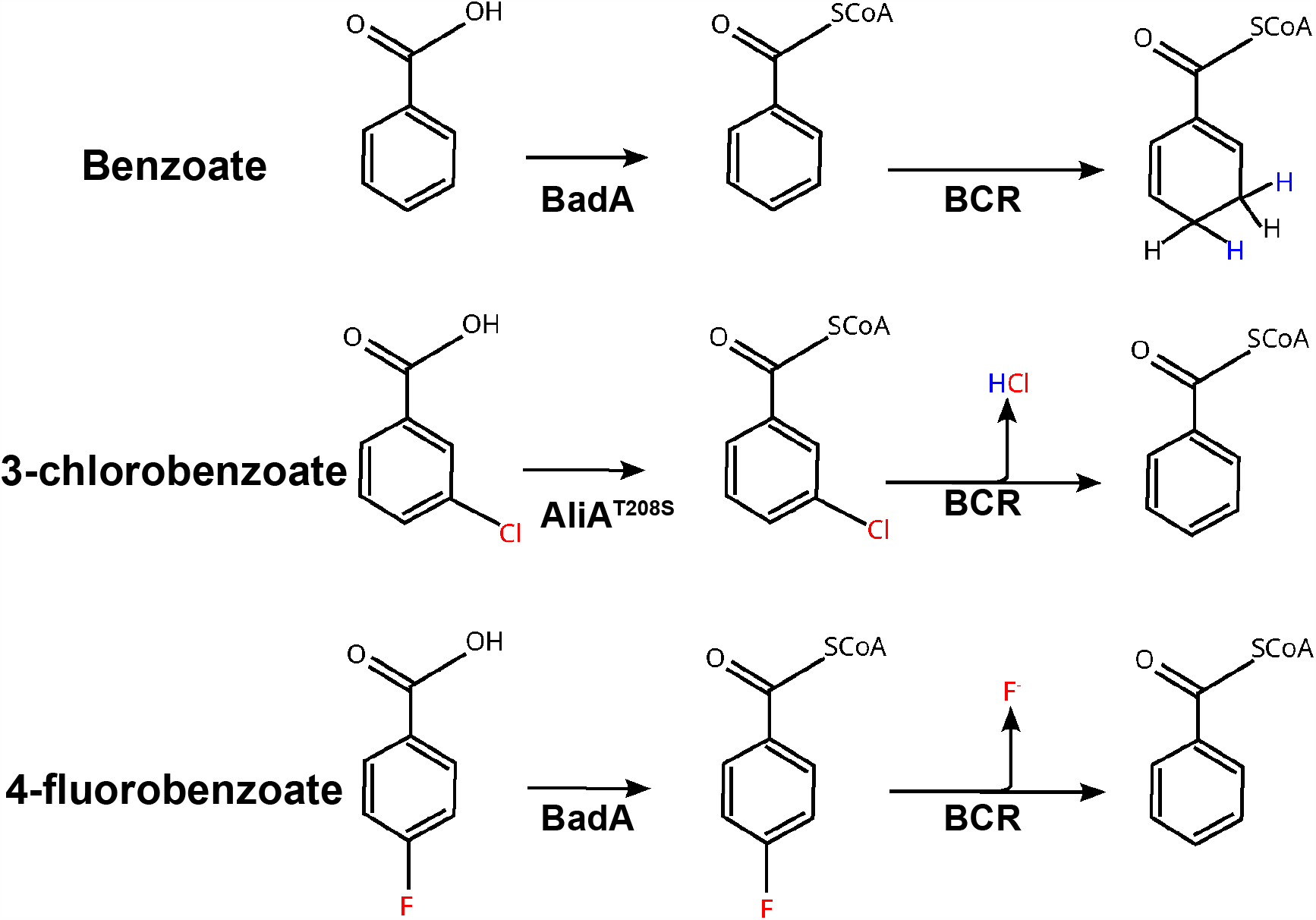
Proposed pathways for benzoate, 3-CBA, and 4-fluoroacetate degradation in *R. palustris*. Although not shown, the 1,5-dienoyl-CoA formed by benzoyl-CoA reductase (BCR) is further metabolized to acetyl-CoA. Both 3-CBA and 4-flourobenzoate generate benzoyl-CoA after dehalogenation, which is then metabolized as shown in the top reaction.

Most strains of *R. palustris*, including the well-known strain CGA009, cannot use 3-CBA as a primary carbon source (9, 10). However, it has been observed that many of these strains can acquire the ability to metabolize 3-CBA within a relatively short period of time (11). This indicates that only one or a few genetic mutations may be necessary to enable 3-CBA metabolism. Presumably, any *R. palustris* strain containing the benzoyl-CoA degradation pathway would be capable of using 3-CBA if the *aliA* allele from RCB100 is introduced. However, introduction of the *aliA*^T208S^ allele in *R. palustris* CGA009 is not sufficient to confer the ability to use 3-CBA as a sole carbon source [(5) and Fig. S1]. This suggests the presence of another unidentified factor that is required for 3-CBA metabolism in *R. palustris*.

To determine what this factor may be, we wanted to reconstruct the evolutionary events that enabled *R. palustris* RCB100 to degrade 3-CBA. To do this, a closely related strain, *R. palustris* CGA009, was experimentally evolved to grow photoheterotrophically using 3-CBA as a sole source of carbon. Whole genome sequencing of the evolved strain and RCB100 led to the discovery of a large deletion that included *badM*, a transcriptional repressor of genes involved in anaerobic aromatic compound degradation. We found that deletion of *badM* is sufficient to allow *R. palustris* CGA009 to degrade 3-CBA and another halogenated aromatic compound, 4-fluorobenzoate. However, *R. palustris* CGA009 *ΔbadM* grew much more slowly than *R. palustris* RCB100 when provided with 3-CBA. Introduction of the *aliA*^T208S^ allele from *R. palustris* RCB100 into *R. palustris* CGA009 Δ*badM* results in a strain with a faster growth rate on 3-CBA. We present a model in which RCB100 initially acquired a deletion that led to constitutive expression of a benzoyl-CoA ligase and benzoyl-CoA reductase and then accumulated other mutations that enhanced its ability to metabolize 3-CBA.

## Results

### Strains capable of degrading 3-CBA contain large deletions near the bad operon

To understand what other factors may be required to enable *R. palustris* to use 3-CBA, we wanted to evolve a strain of *R. palustris* to use 3-CBA as its sole carbon source during photoheterotrophic growth. We wanted to find a closely related strain unable to metabolize 3-CBA for use in adaptive evolution so that comparisons could be drawn with *R. palustris* RCB100, allowing reconstruction of how RCB100 evolved to metabolize 3-CBA. We compared the complete genome sequences of 13 *R. palustris* strains by analyzing average nucleotide identity (ANI) with BLASTn, which revealed that strain CGA009 is the most closely related *R. palustris* strain to RCB100 with an ANI score of 0.99, corresponding to 99% identity between the two genomes (Fig. 2A). Both *R. palustris* CGA009 and *R. palustris* RCB100 were isolated near Ithaca, New York, which is consistent with these two strains being the most closely related (3, 12).

**Figure 2.**
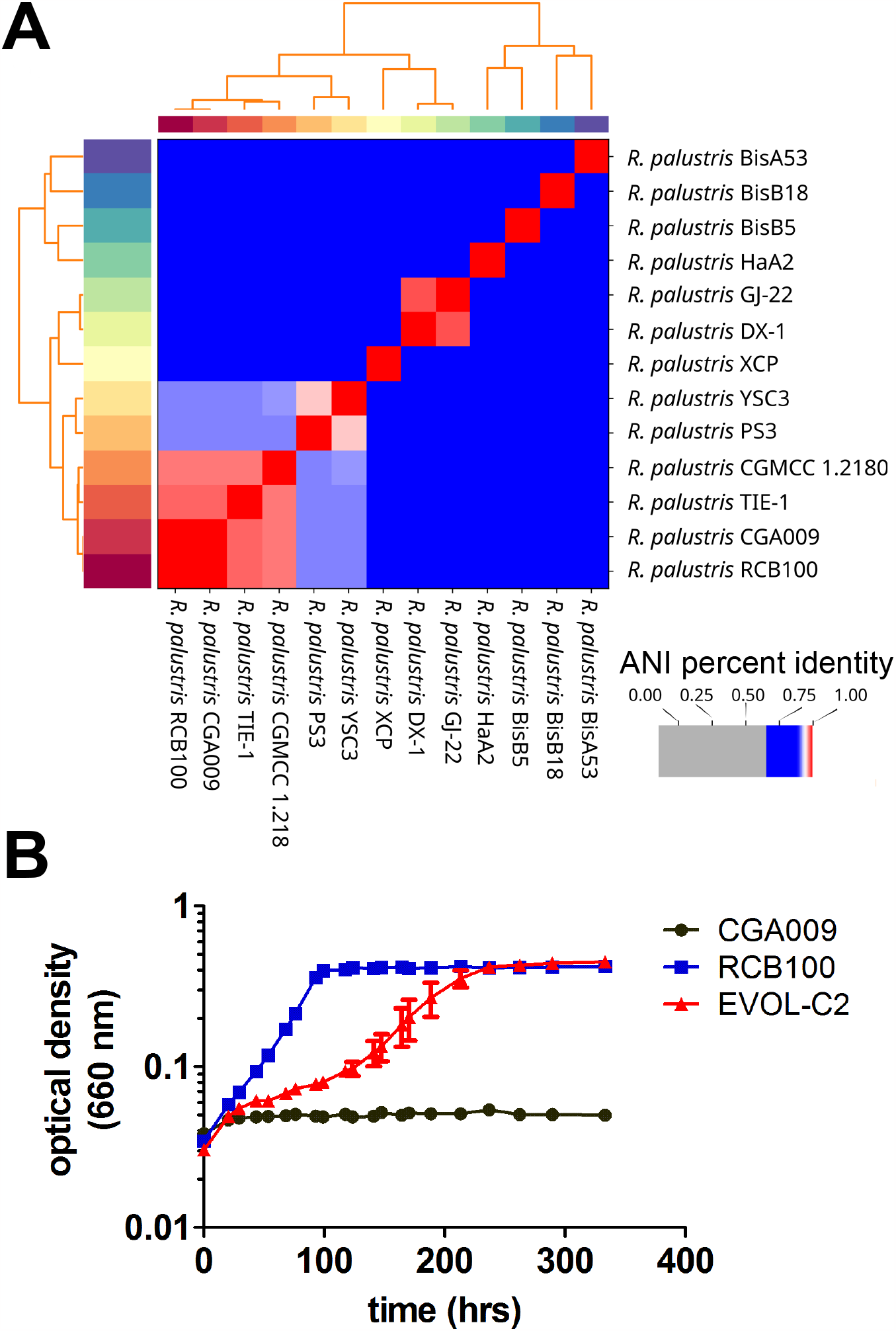
Isolation of a strain of *R. palustris* CGA009 capable of growing with 3-CBA as its sole carbon source. (A) As shown by average nucleotide identity (ANI), the genomes of the closely-related *R. palustris* RCB100 and *R. palustris* CGA009 are >95% identical. (B) Growth of *R. palustris* CGA009, *R. palustris* RCB100, and a strain of *R. palustris* CGA009 adapted to grow with 3-CBA (EVOL-C2) in minimal medium with 1 mM 3-CBA. Data are from three independent cultures and error bars represent standard deviation.

In addition to being closely related to RCB100, *R. palustris* CGA009 is unable to grow with 3-CBA as a carbon source, and we confirmed that introduction of *aliA*^T208S^ is not sufficient to enable *R. palustris* CGA009 to grow with 3-CBA as its sole carbon source (Fig. S1) (5). To select for an adapted strain that can grow with 3-CBA as the sole carbon source, CGA009 was grown in medium with a mixture of benzoate and 3-CBA for several months in the light. After three successive transfers of enrichment cultures into liquid medium with an increasing ratio of 3-CBA to benzoate, we isolated an evolved strain (EVOL-C2) of *R. palustris* that was able to grow with 3-CBA as the sole carbon source. Unlike strain CGA009, EVOL-C2 can use 3-CBA as a sole source of carbon, although it grew much more slowly than strain RCB100 (Fig. 2B).

To determine the genetic basis of the evolved phenotype, we sequenced the genome of strain EVOL-C2 and RCB100 (13) and compared its genome to strain CGA009. Unexpectedly, we found that both EVOL-C2 and RCB100 contained large deletions near the *bad* operon, which encodes genes required benzoyl-CoA degradation (Fig. 3A, Table S1). In EVOL-C2, the deleted region (6,521 bp) included partial deletion of *badM* and *hbaA*, and complete deletion of *badL, hbaH, hbaG, hbaF* and *hbaE*. RCB100 also contained a similar but significantly larger (10,387 bp) deletion that encompasses *badM* through *hbaC* (Fig. 3A, Table S1). This deletion was the only genetic change that was shared between EVOL-C2 and RCB100 (Table S1).

**Figure 3.**
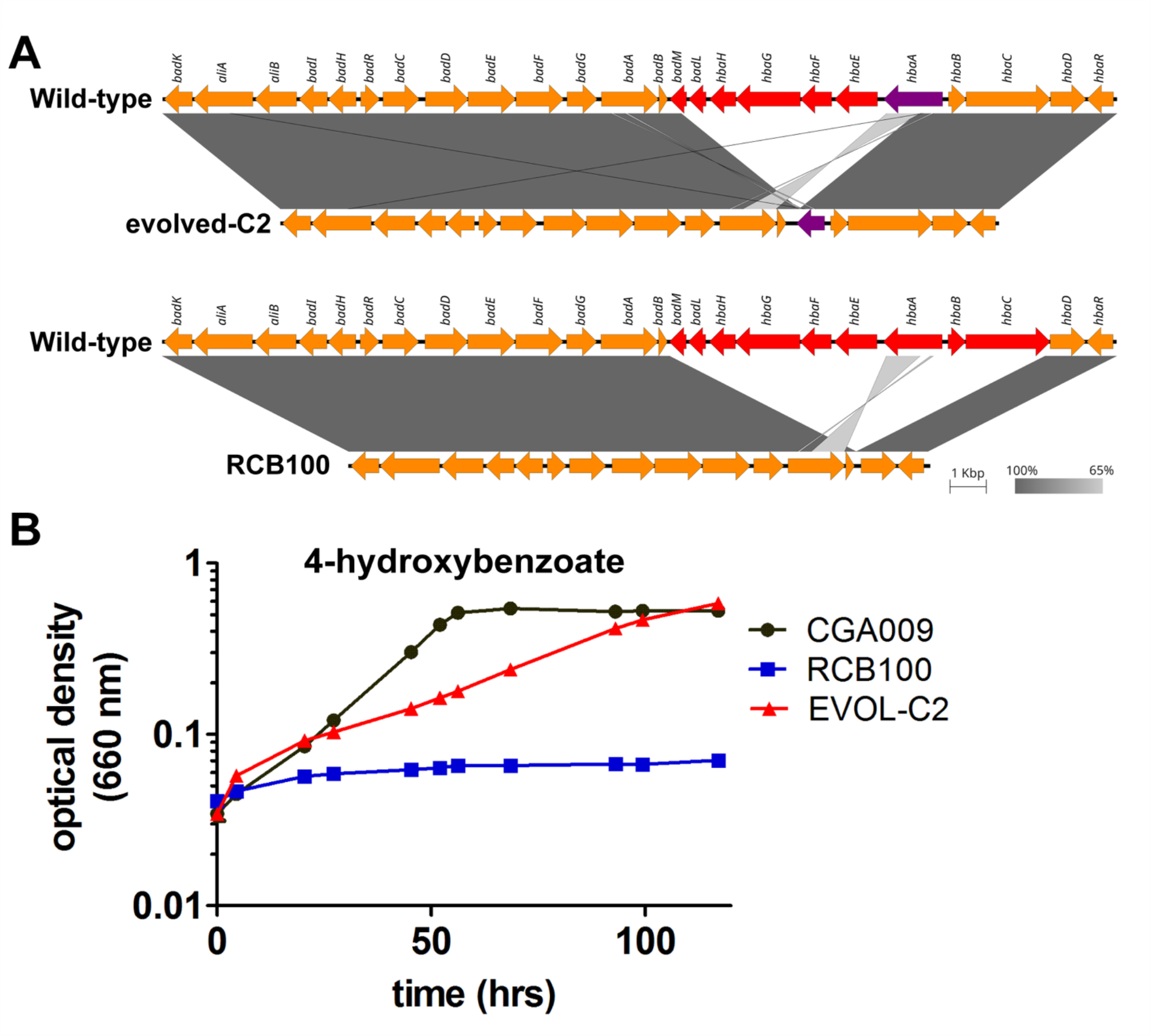
*R. palustris* RCB100 and *R. palustris* EVOL-C2 have large deletions near the *bad* operon. (A) represents comparison of strains CGA009 and EVOL-C2 (top) and comparison of CGA009 with RCB100 (bottom). Arrows indicate genes and their direction with the gene names listed on top of the arrows. The deleted genes are shown in red whereas partially deleted genes are shown in purple. The shaded gray gradient represents the degree of homology between the compared regions. The plots were generated with Easyfig using the BLASTn algorithm. (B) Growth of *R. palustris* CGA009, *R. palustris* RCB100, and *R. palustris* CGA009 EVOL-C2 in minimal medium with 6 mM 4-HBA. Data are from three independent cultures and error bars represent standard deviation.

The *hba* genes are associated with degradation of 4-hydroxybenzoate (4-HBA). Genes *hbaHGFE* encode a predicted ABC transport system for 4-HBA (4). Like benzoate, 4-HBA is activated by a 4-HBA-CoA ligase encoded by *hbaA*, and *hbaBC* encode components of 4-HBA-CoA reductase that reductively removes the hydroxyl group from 4-HBA-CoA to yield benzoyl-CoA. Deletion of these genes would be expected to impact the ability of EVOL-C2 and RCB100 to grow with 4-HBA as the sole carbon source. As shown in Fig. 3B, EVOL-C2 grew more slowly on 4-HBA compared to its parent strain, and RCB100 was unable to grow with 4-HBA as a sole carbon source, confirming the presence of deletions that disrupt 4-HBA metabolism.

### Deletion of the transcriptional repressor badM in R. palustris CGA009 is sufficient to enable degradation of 3-CBA

Two additional genes, *badL* and *badM*, were also deleted in EVOL-C2 and RCB100 (Fig. 3A). BadL and BadM play a role in regulation of the *bad* operon, *badDEFGAB*. The *bad* operon encodes the benzoyl-CoA ligase, BadA; a ferredoxin, BadB; and the subunits for benzoyl-CoA reductase, BadDEFG. BadM is a transcriptional repressor of the *bad* operon, and BadM bound to acetamidobenzoates leads to derepression of *bad* operon expression (14–16). In a *badM* mutant, the *bad* operon is constitutively expressed (15). We hypothesized that constitutive expression of the *bad* operon in EVOL-C2 and RCB100 may be sufficient to enable these strains to use 3-CBA as a sole carbon source. We tested the ability of *R. palustris* CGA009 Δ*badM* to grow photoheterotrophically with 3-CBA as a carbon source. As shown in Fig. 4A and 4B, wild-type *R. palustris* CGA009 and *R. palustris* CGA009 Δ*badM* grow at similar rates when benzoate is provided as a carbon source, but only *R. palustris* CGA009 Δ*badM* can grow when 3-CBA is provided as a carbon source. This indicates that constitutive expression of the *bad* operon is sufficient for *R. palustris* CGA009 to grow on 3-CBA. However, *R. palustris* CGA009 Δ*badM* still grew slower than RCB100 or EVOL-C2.

**Figure 4.**
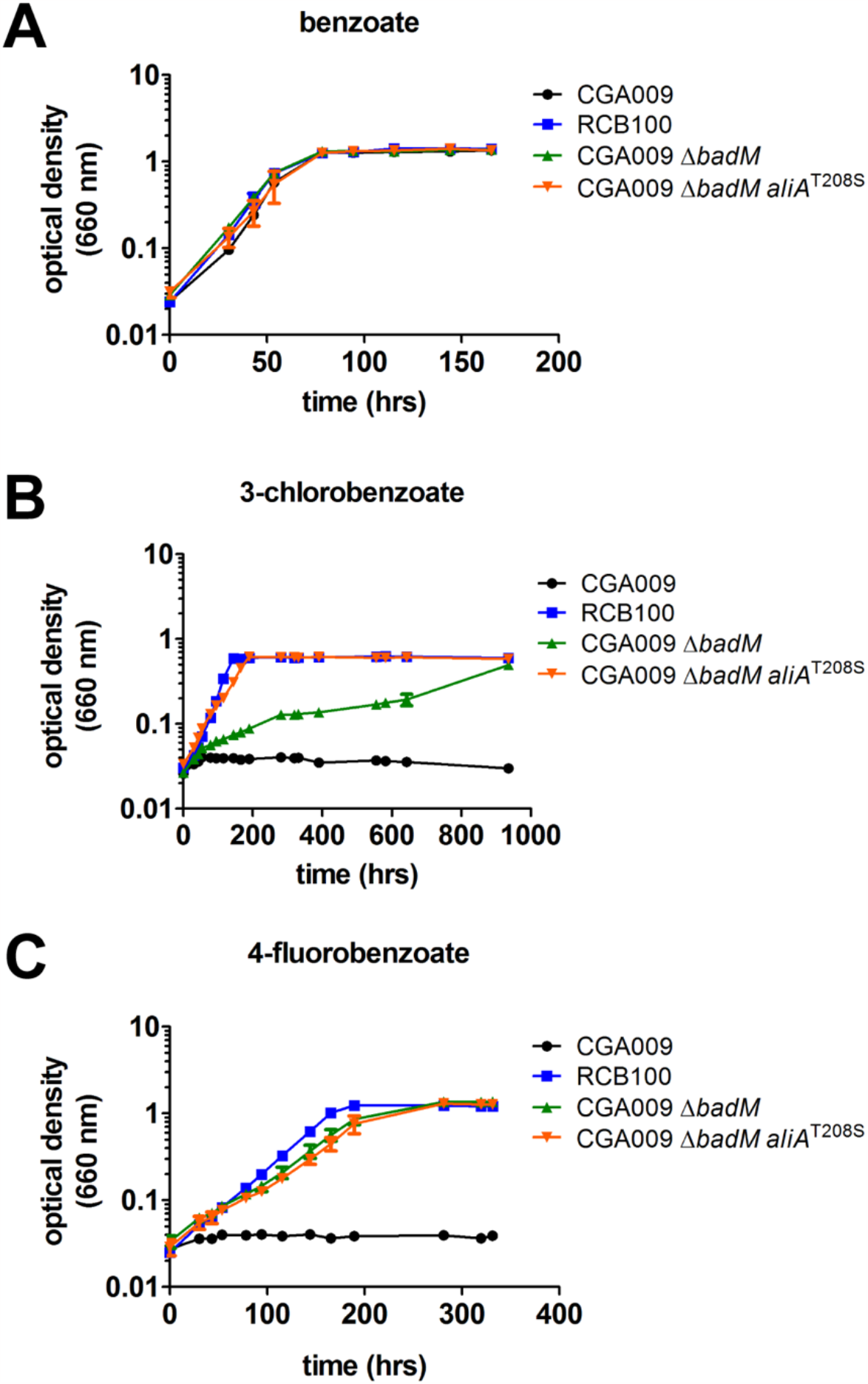
Deletion of *badM* enables growth of *R. palustris* on 3-CBA and 4-fluorobenzoate, and *aliA*^T208S^ enhances growth of a *badM* mutant on 3-CBA. Growth of *R. palustris* strains CGA009, RCB100, CGA009 with a deletion in *badM* (CGA009 Δ*badM*), and CGA009 Δ*badM* encoding the allele from RCB100 (CGA009 Δ*badM aliA*^T208S^) in (A) minimal medium supplemented with 5.7 mM benzoate; (B) minimal medium supplemented with 3 mM 3-CBA; and (C) minimal medium supplemented with 5 mM 4-fluorobenzoate. Data are from three independent cultures and error bars represent standard deviation.

EVOL-C2 also contained a single mutation in the transcription regulator *badR*, which results in a protein variant with arginine 80 substituted for a leucine. BadR represses expression of the cyclohexanecarboxylate (CHC) operon, which includes *aliA* (*badHI-aliAB-badK)* (17). Since a variant of AliA plays a role in 3-CBA metabolism in RCB100, we wanted to determine if inactivation of *badR* plays a role in 3-CBA metabolism. To test this, *R. palustris* CGA009 Δ*badM, R. palustris* CGA009 Δ*badR*, and a *R. palustris* CGA009 Δ*badM badR* were grown photoheterotrophically with 3-CBA as a carbon source. We found that *R. palustris* CGA009 Δ*badM* and *R. palustris* CGA009 Δ*badM badR* grew at similar rates on 3-CBA (Fig. S2). However, *R. palustris* CGA009 Δ*badR* was unable to grow on 3-CBA as a sole source of carbon (Fig. S2). This indicates that deletion of *badR* even when in combination with a deletion in *badM* does not enhance degradation of 3-CBA by *R. palustris* CGA009.

We next wanted to determine if deletion of *badM* was sufficient to enable growth on other halogenated aromatic compounds. Unlike 3-CBA, which requires a CoA ligase that has more activity with 3-CBA, 4-fluorobenzoate can be activated by benzoyl-CoA ligase at similar rates to benzoate, and the C-F bond can be reductively cleaved by benzoyl-CoA reductase to form benzoate (Fig. 1) (8). We hypothesized that deletion of *badM* in *R. palustris* CGA009 would enable it to use 4-fluorobenzoate as a carbon source. As shown in Fig. 4C, *R. palustris* CGA009 is unable to use 4-fluorobenzoate as growth substrate, but *R. palustris* CGA009 Δ*badM* and RCB100 were able to grow. This indicates that deletion of *badM* enables CGA009 to use other halogenated aromatic compounds.

### Introduction of aliA^T208S^ in R. palustris CGA009 ΔbadM improves growth with 3-CBA but not 4-fluorobenzoate

Although *R. palustris* CGA009 Δ*badM* can grow when 3-CBA is provided, it grows slowly compared to RCB100 (Fig. 4B). This is likely because, in addition to constitutive expression of benzoyl-CoA reductase, RCB100 also expresses a CoA ligase (AliA^T208S^) that has 10 times more activity with 3-CBA than the AliA from CGA009 (5). To test this, the *aliA*^T208S^ allele was introduced into *R. palustris* CGA009 Δ*badM*. As shown in Fig. 4A, there is no difference in growth between these strains when they are provided with benzoate as a carbon source. However, when provided with 3-CBA as a carbon source, *R. palustris* CGA009 Δ*badM aliA*^*T208S*^ cells grew significantly faster than *R. palustris* CGA009 Δ*badM* (Fig. 4B). Both *R. palustris* CGA009 Δ*badM* and *R. palustris* CGA009 Δ*badM aliA*^*T208S*^ grew at similar rates on 4-fluorobenzoate, confirming that AliA^T208S^ is only required for 3-CBA degradation (Fig. 4C). This indicates that deletion of *badM* and *aliA*^T208S^ is sufficient to reproduce the ability of RCB100 to degrade 3-CBA in another *R. palustris* strain.

## Discussion

Using adaptive laboratory evolution, we reconstructed the evolutionary events that enabled a natural isolate to use halogenated aromatic compounds. We found that deletion of the transcriptional repressor, *badM*, is sufficient to enable *R. palustris* to use 3-CBA as a carbon source, indicating that derepression of the *bad* genes is required for *R. palustris* to degrade 3-CBA. This is consistent with the observation that *R. palustris* strains can co-metabolize 3-CBA and benzoate despite being unable to metabolize 3-CBA as the sole source of carbon (3, 9, 10). We also found that introduction of *aliA*^T208S^ into *R. palustris* CGA009 was not sufficient to enable growth on 3-CBA, but introduction of *aliA*^T208S^ into *R. palustris* CGA009 Δ*badM* significantly increased its growth rate on 3-CBA. This indicates that RCB100 likely evolved by first acquiring a deletion that inactivated *badM*, leading to constitutive expression of a CoA ligase and benzoyl-CoA reductase. However, the rate-limiting step in this strain must be CoA activation, leading to a strong selective pressure to acquire a mutation that improved CoA ligase activity with 3-CBA.

The fact that *R. palustris* CGA009 Δ*badM* can grow on 3-CBA indicates that native CoA ligases have activity against 3-CBA. This is surprising because the ability to degrade 3-CBA is thought to be restricted to bacterial strains that encode a CoA ligase that has increased activity with 3-CBA (6, 18). *R. palustris* CGA009 and RCB100 encode three CoA ligases involved in anaerobic aromatic compound degradation: benzoyl-CoA ligase, BadA 4-hydroxybenzoyl-CoA ligase, HbaA; and cyclohexanecarboxylate-CoA ligase, AliA. Wild-type AliA is unlikely to be involved since it is constitutively expressed in a *badR* mutant. *R. palustris* Δ*badR* is unable to grow on 3-CBA and deletion of both *badR* and *badM* does not significantly improve growth on 3-CBA. HbaA is also not involved since *hbaA* is deleted in EVOL-C2 and RCB100. However, deletion of *badM* leads to constitutive expression of BadA (15). This implicates BadA in degradation of 3-CBA. BadA has been shown to have some activity against 3-CBA *in vitro*, and BadA activity is even higher in a *badM* mutant compared to induced wild-type cells (15, 19). This suggests that there is enough promiscuous activity by BadA to enable growth on 3-CBA, and a CoA ligase specific to 3-CBA is not a requirement *per se*.

Our findings also indicate that *R. palustris* is unable to sense halogenated aromatic compounds since deletion of *badM* is required for *R. palustris* to grow on these compounds. Repression of BadM is relieved when it is bound to acetaminobenzoates, which are produced by the acetylation of aminobenzoates by BadL (14). When grown with 3-CBA or 4-fluorobenzoate, accumulation of acetaminobenzoates must not occur in *R. palustris*, and BadM repression is not relieved. It is difficult to speculate why BadM repression is not relieved in the presence of halogenated compounds as it is still unclear why acetaminobenzoates accumulate when cells are grown with benzoate. The growth medium for *R. palustris* contains 4-aminobenzoate (11 μM) for folate biosynthesis, and 4-aminobenzoate is a substrate for BadL (14, 20). Why derepression of BadM only occurs in the presence of aromatic compounds like benzoate but not in the presence of halogenated aromatic compounds in medium containing 4-aminobenzoate is unclear.

This is the first evolutionary reconstruction of an environmental isolate capable of using halogenated aromatic compounds. Adaptive laboratory evolution has provided numerous examples of how regulatory mutations or variants of a promiscuous enzyme can enable degradation of man-made compounds (21– 29). Here, we show that both bottlenecks, gene regulation and substrate specificity of a promiscuous enzyme, were overcome to enable efficient degradation of 3-CBA. However, there appears to be an ecological trade-off associated with the ability to degrade 3-CBA as both EVOL-C2 and RCB100 have a reduced ability to grow on 4-HBA. Specialization in consuming an available resource at the expense of losing the ability to consume other resources can occur when bacterial populations are switched from a heterogenous environment to a more homogenous environment (30). This suggests that *R. palustris* RCB100 likely evolved in an environment rich in 3-CBA but lacking other aromatic compounds like 4-HBA. While the presence of natural substrates could enable co-metabolism of a man-made compound, it could also relax selection and delay evolution of microorganisms that can metabolize the compound on its own (31). This could confound our ability to identify environmental isolates capable of degrading man-made compounds, particularly compounds only recently released into the environment.

## Material and Methods

### Bacterial strains and culture conditions

All bacterial strains used in the current study are listed in Table S2. *R. palustris* strains were grown under anoxic conditions in 10 mL of minimal mineral medium (PM) in sealed Hungate tubes (12, 20). Carbon substrates from anoxic, sterile stock solutions were added after autoclaving. Carbon sources were added to the following final concentrations where indicated: 10 mM succinate, 20 mM acetate, 5.7 mM benzoate, 1 to 3 mM 3-chlorobenzoate, 5 mM 4-fluorobenzoate, and 6 mM 4-HBA. All liquid cultures were grown at 30°C and provided light (30 μmol photons/m^2^/s) from a 60W halogen light bulb (General Electric). Cultures were initially grown in PM with succinate and then diluted in PM with the stated carbon source for growth analysis. Optical density was determined at 660 nm using a Genesys 50 UV-visible spectrophotometer. *Escherichia coli* strains were grown in lysogeny broth (LB) medium at 37°C. When necessary, *R. palustris* and *E. coli* cultures were supplemented with gentamicin (Gm) at concentrations of 100 and 20 μg/mL, respectively.

### *Experimental evolution of* R. palustris *CGA009 and isolation of evolved* R. palustris *EVOL-C2*

A single colony of *R. palustris* CGA009 was picked up with a sterilized toothpick and resuspended in 200 μL of PM medium in a sterile eppendorf tube before the suspended cells were introduced into sealed Hungate tubes containing 10 mL of PM supplemented with acetate. To adapt *R. palustris* CGA009 to grow on 3-CBA, cells were inoculated into 4 tubes containing PM supplemented with 5.7 mM benzoate and 1 mM 3-CBA as described in (11). Cells were grown for three weeks under light, before they were transferred to PM medium supplemented with 2.85 mM benzoate and 1.5 mM 3-CBA. Cells were grown in the light for five weeks and then transferred into PM medium supplemented with 1.5 mM benzoate and 2.5 mM 3-CBA. Cells were grown in the light for five weeks before they were transferred to PM medium with 0.5 mM benzoate and 3.5 mM 3-CBA. Cells were grown in the light for five weeks and transferred into PM supplemented with 3.5 mM 3-CBA. After nine weeks, a single replicate grew, and 100 μL of cells were spread on plates containing PM with 5.7 mM benzoate. After three weeks incubation, 13 random colonies were selected and grown in hungate tubes with PM supplemented with 1 mM 3-CBA. EVOL-C2 was one of the 13 colonies that consistently grew in PM supplemented with 1 mM 3-CBA. After repeated confirmation of growth on liquid PM supplemented with 1 mM 3-CBA, 100 μL of EVOL-C2 was transferred to PM liquid medium supplemented with 20 mM acetate and 0.1% yeast extract. Once fully grown, a glycerol stock was prepared and stored at –80°C and genomic DNA was extracted for sequencing as described below.

### Genomic DNA extraction and whole genome sequence analyses

Genomic DNA was extracted as described previously (32). Briefly, *R. palustris* EVOL-C2 was grown in 10 mL PM medium with 20 mL acetate. Freshly grown *R. palustris* cells were harvested with centrifugation at 12,000 rpm for 10 minutes and genomic DNA was extracted from cell pellets as described (6, 8). Both Illumina and Nanopore libraries were prepared at Microbial Genome Sequencing Center (MiGS; https://www.migscenter.com/), using the workflows reported previously (6, 9), before sequencing on NextSeq 550 and MinION v9.4.1 flow cell, respectively. Base calling was performed using guppy v.4.2.2 with default parameters *+ effbaf8*. The quality control and adapter trimming for Illumina reads was performed with bcl2fastq v.2.20.0.445 (10), whereas porechop v.0.2.3_seqan2.1.1 (11) was used for Oxford Nanopore Technology (ONT) sequencing reads. The Illumina dataset comprised of 5,063,564 reads pairs and 1,632,037,238 bases whereas Nanopore sequencing produced 116,959 reads and 303,624,697 bases.

A hybrid genome assembly was generated from Illumina and ONT reads with Unicycler v.0.4.8 (12) and QUAST v.5.0.2 (13) was used to obtain genome assembly statistics. Genome assembly resulted in two circular contigs of 5,452,700 bp and 8,427 bp that represented chromosome and plasmid, respectively. The GC (%) of the assemble genome was 65.03, whereas the length of the genome was 5,461,127 bp. The Assembly annotation was performed with NCBI Prokaryotic Genome Annotation Pipeline (PGAP). To predict mutations in strains EVOL-C2 and RCB100 in comparison to strain CGA009, we used breseq v0.36.0 (33).

We used PYANI v0.2.10 [(34); https://github.com/widdowquinn/pyani] to calculate average nucleotide identity among 13 selected *R. palustris* genomes based on BLASTN+ (ANIb) method. Easyfig v2.2.2 (35) was used to generate genome comparison visualization plots.

### *Genetic manipulation of* R. palustris

A construct containing the *aliA* locus from *R. palustris* RCB100 was constructed for allelic exchange using the primers shown in Table S2. Genomic DNA purified from *R. palustris* RCB100 was used as a template for PCR amplification using Phusion High-Fidelity DNA polymerase(New England Biolabs) with primers *aliA*F and *badK*R (Table S2). Using homologous recombination as described in (36), the resulting product was incorporated into linearized pJQ200SK generated from PCR amplification with primers pJQ200SK-*aliA*F and pJQ200SK-*badK*R (Table S2). The resulting plasmid was mobilized into *R. palustris* CGA009 by conjugation with *E. coli* S17-1 and double-crossover events for allelic exchange was accomplished using a previously described selection and screening strategy (37). Replacement of *aliA* in *R. palustris* CGA009 with the *aliA* locus from *R. palustris* RCB100 was verified using PCR followed by Sanger sequencing (Azenta Life Sciences, Burlington, MA).

## Data availability

The *R. palustris* RCB100 genome sequence is available at NCBI under the accession number <u>GCF_016584445.1</u>. The *R. palustris* CGA009 genome sequence is available under the accession number <u>GCF_000195775.2</u>, and the *R. palustris* CGA009 EVOL-C2 genome sequence is available under the accession number <u>GCA_031600295.1</u>. Raw sequencing data for EVOL-C2 is available at NCBI under the accession numbers SRR17562209 (Oxford nanopore) and SRR17562208 (Illumina). The accession number for the EVOL-C2 sequencing BioProject is PRJNA796295, and the Biosample accession number is SAMN24839554.

## Acknowledgments

We thank Caroline Harwood and Yasuhiro Oda for providing mutant strains of *R. palustris* used in this study. We also thank Nathan Lewis, Nicholas Haas and Jack Reddan for valuable discussions during the course of this study. This work was supported through the Biocatalysis Initiative grant awarded to K.R.F. by the BioTechnology Institute, University of Minnesota.

